# Quorum sensing regulation of a major facilitator superfamily transporter affects multiple streptococcal virulence factors

**DOI:** 10.1101/2022.05.06.490991

**Authors:** Jennifer C. Chang, Reid V. Wilkening, Kate M. Rahbari, Michael J. Federle

## Abstract

Cell-cell signaling mediated by Rgg-family transcription factors and their cognate pheromones is conserved in *Firmicutes*, including all streptococci. In *Streptococcus pyogenes*, or Group A strep, one of these systems, the Rgg2/3 quorum sensing (QS) system, has been shown to regulate phenotypes including cellular aggregation and biofilm formation, lysozyme resistance, and macrophage immunosuppression. Here, we show that the abundance of several secreted virulence factors (streptolysin O, SpyCEP, and M protein) decreases upon induction of QS. The main mechanism underlying the changes in protein levels appears to be transcriptional, occurs downstream of the QS circuit and is dysregulated by the deletion of an Rgg2/3 QS-regulated major facilitator superfamily (MFS) transporter. Additionally, we identify this MFS transporter as the factor responsible for a previously observed increase in aminoglycoside sensitivity in QS-induced cells.

**Importance:** The production of virulence factors is a tightly regulated process in bacterial pathogens. Efforts to elucidate the mechanisms by which genes are regulated may advance the understanding of factors influencing pathogen behavior or cellular physiology. This work finds that expression of a major facilitator superfamily (MFS) transporter, which is governed by a quorum sensing (QS) system, impacts the expression of multiple secreted virulence factors and accounts for a documented QS-dependent antibiotic susceptibility. Although the mechanism underlying this effect is not clear, MFS orthologs with high sequence similarity from *S. pneumoniae* and *S. porcinus* were unable to substitute indicating substrate specificity of the GAS MFS gene. These findings demonstrate novel associations between the expression of a transmembrane transporter and virulence factor expression and aminoglycoside transport.

## Introduction

*Streptococcus pyogenes* (Group A Streptococcus; GAS) is a gram-positive obligate human pathogen that may asymptomatically colonize an individual or cause disease ranging from mild (strep throat, impetigo) to severe (bacteremia, necrotizing fasciitis). A serious post-infection complication, rheumatic heart disease, accounts for the greatest global GAS disease burden with an estimated 33 million people affected by this condition resulting in more than 300,000 deaths per year ^1^. Despite decades of efforts, a vaccine to prevent GAS infection has remained elusive.

Bacteria may seem simple compared to multicellular organisms, but they possess multiple systems, including some that can influence group behavior, that allow them to sense and respond to minute environmental changes. Quorum sensing is a phenomenon in which bacterial cells produce and respond to small signaling molecules in a concentration-dependent manner, resulting in the population-wide coordination of behaviors such as biofilm formation, virulence factor production, or natural transformation. These signaling molecules, called pheromones, can be intercepted by different species, thus allowing for both intercellular and interspecies communication.

Four types of quorum-sensing systems have been described for GAS: Sil, lantibiotic systems, LuxS/AI-2, and regulator gene of glucosyltransferase (Rgg) systems

^2^. Of the latter, GAS possess four Rgg-type transcription factors which interact with small oligopeptide pheromones to control three discrete regulons: RopB (Rgg1) mediates expression of the SpeB cysteine protease, a well-characterized virulence factor; ComR (Rgg4) is implicated in natural competence; and Rgg2 and Rgg3 together regulate genes linked to cell aggregation, biofilm formation, lysozyme resistance, nasopharyngeal colonization, pharyngitis and macrophage immunosuppression^3-10^. The Rgg2/3 system is unique in that the two homologous regulators recognize the same pheromones (<u>s</u>hort <u>h</u>ydrophobic <u>p</u>eptides 2 and 3; SHP2 and SHP3) yet exert opposing effects on the same gene targets. Rgg2, the positive transcriptional regulator, is required for target gene expression and quorum-sensing induction; in contrast, Rgg3 is a transcriptional repressor, and its deletion results in constitutive auto-induction or a quorum-sensing locked ON phenotype.

GAS produce a wide array of virulence factors that contribute to pathogenesis. Given the cell surface changes we previously observed in response to Rgg2/3 QS, we expanded our survey to include other cell surface-associated virulence factors. While we found no evidence of direct regulation by Rgg2 or Rgg3, we demonstrate that the expression of at least three secreted factors, streptolysin O, SpyCEP, and M protein, is indirectly regulated by Rgg2/3 quorum sensing through its control of a major facilitator superfamily transporter, Spy49_0460.

## Results

### Rgg2/3 signaling decreases hemolytic activity attributable to streptolysin O

Previous studies of the Rgg2/3 signaling pathway suggest that expression of this regulon results in changes to the cell surface, resulting in phenotypes including increased aggregation, biofilm formation and lysozyme resistance ^3, 4^. To test whether Rgg-SHP signaling altered additional cell surface-associated or secreted virulence factors of GAS, bacteria were treated with the activating peptide SHP3-C8 (C8) or the nonfunctional reverse C8 sequence (rev) to achieve “quorum sensing ON” (QS-ON) and QS-OFF states, respectively. Culture supernatants were collected at late logarithmic phase, and hemolytic activity was determined using a red blood cell lysis assay. GAS produces two hemolysins that contribute to its β-hemolytic phenotype on blood agar, an oxygen-stable bacteriocin called streptolysin S (SLS) which belongs to the family of thiazole/oxazole-modified microcins ^11^ and a cholesterol-dependent, pore-forming, oxygen-labile toxin, streptolysin O (SLO) ^12^. Assessment of Δ*sagA* (SLS^-^) and Δ*slo* mutants, as well as selective inhibition of SLO by the addition of cholesterol, demonstrated that the reduction in hemolytic activity in QS-ON cells was due to streptolysin O (Fig. 1A). Subsequent western blot analysis confirmed the reduction of SLO in culture supernatants from C8-treated, QS-ON wild-type cells, and that this response requires the transcriptional activator Rgg2 (Fig. 1B). As expected, supernatants from the Δ*rgg3* mutant, which is QS-ON in CDM even in the absence of induction with synthetic C8, contained less SLO than WT; treatment with synthetic C8 led to a further reduction of SLO. In agreement with our observations from the red blood cell lysis assay, deletion of the SLS structural gene *sagA* had no effect on the reduction of SLO in response to QS induction.

**Figure 1.**
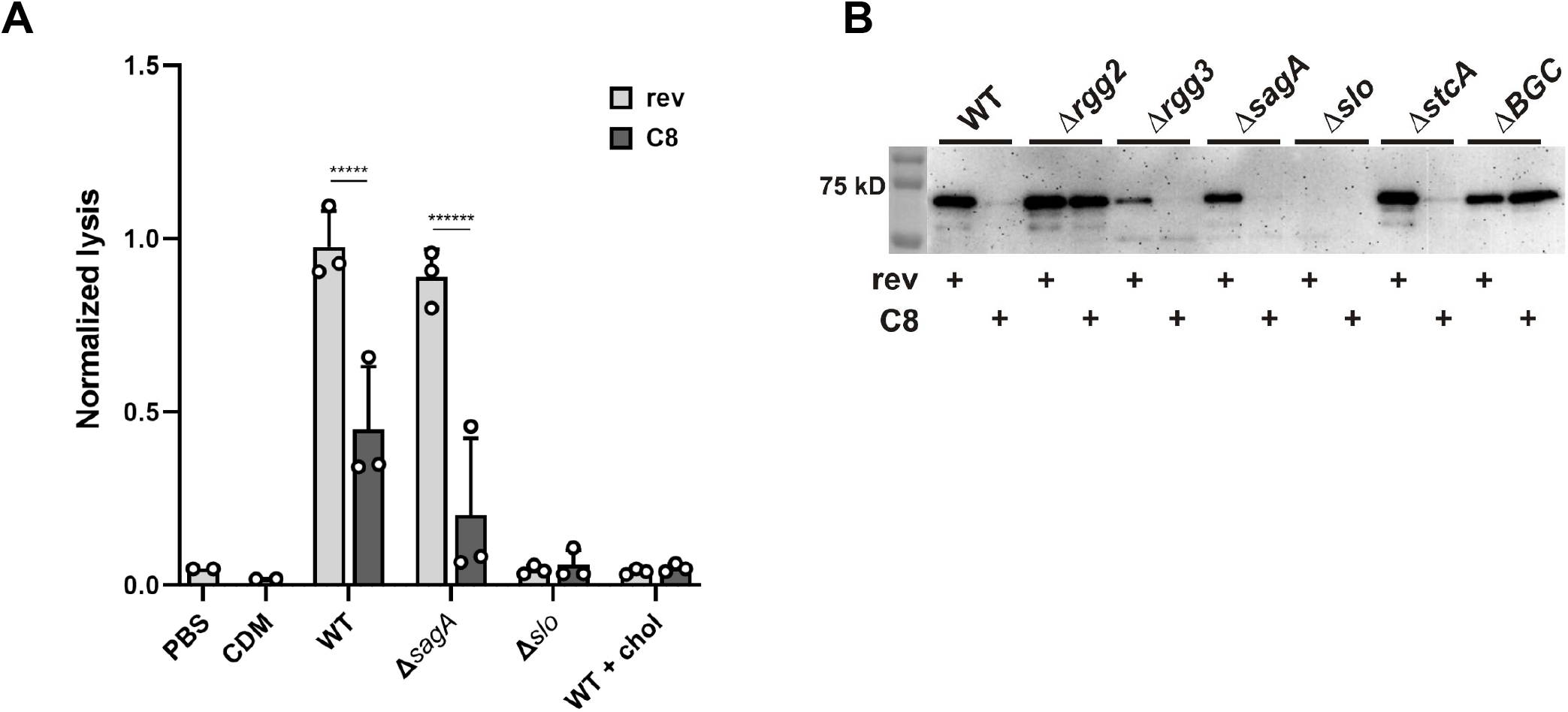
Reduced hemolysis in QS-ON cells is due to streptolysin O. Late log phase supernatants from quorum-sensing off (rev) or quorum-sensing on (C8) bacteria were collected and analyzed for **A**. Hemolytic activity against sheep red blood cells; and **B**. The presence of streptolysin O by western blot. Data shown are the mean and SD of three biological replicates (A) and are representative of at least three experiments (A and B). *****, P=0.00001 and ******, P<0.000001, by unpaired t-test.

### Decrease in SLO is linked to a major facilitator superfamily transporter

Induction of Rgg2/3 signaling leads to transcription at two primary sites in the genome – the genes divergently transcribed from each *rgg*, including its respective *shp* pheromone, and downstream operons^4^(Fig. 2A). Proximal to *rgg2/shp2* is a small, positively-charged protein, StcA, which contributes to QS-induced biofilm formation and resistance to lysozyme ^5^. The genes downstream of *shp3* appear to encode a biosynthetic gene cluster (BGC) and have recently been linked to the suppression of NFkB activation and proinflammatory cytokine responses of macrophages ^6^. Western blotting determined that the decrease in SLO in QS-ON culture supernatants was attributable to genes in the BGC, not *stcA* (Fig. 1B). Using a series of deletion mutants (Fig. 2A), we further determined that the reduction of SLO was dependent solely on a major facilitator superfamily (MFS) protein annotated as a putative efflux protein (*spy49_0460*) encoded at the distal end of the cluster (Fig. 2B). We confirmed that deletion of *spy49_0460* did not affect the ability of this mutant to respond to synthetic peptide or diminish the ability of the *rgg3 spy49_0460* double mutant to auto-induce the signaling system (Fig. S1). These data indicate that the influence of Spy49_0460 on SLO production lies downstream of the QS circuit.

**Figure 2.**
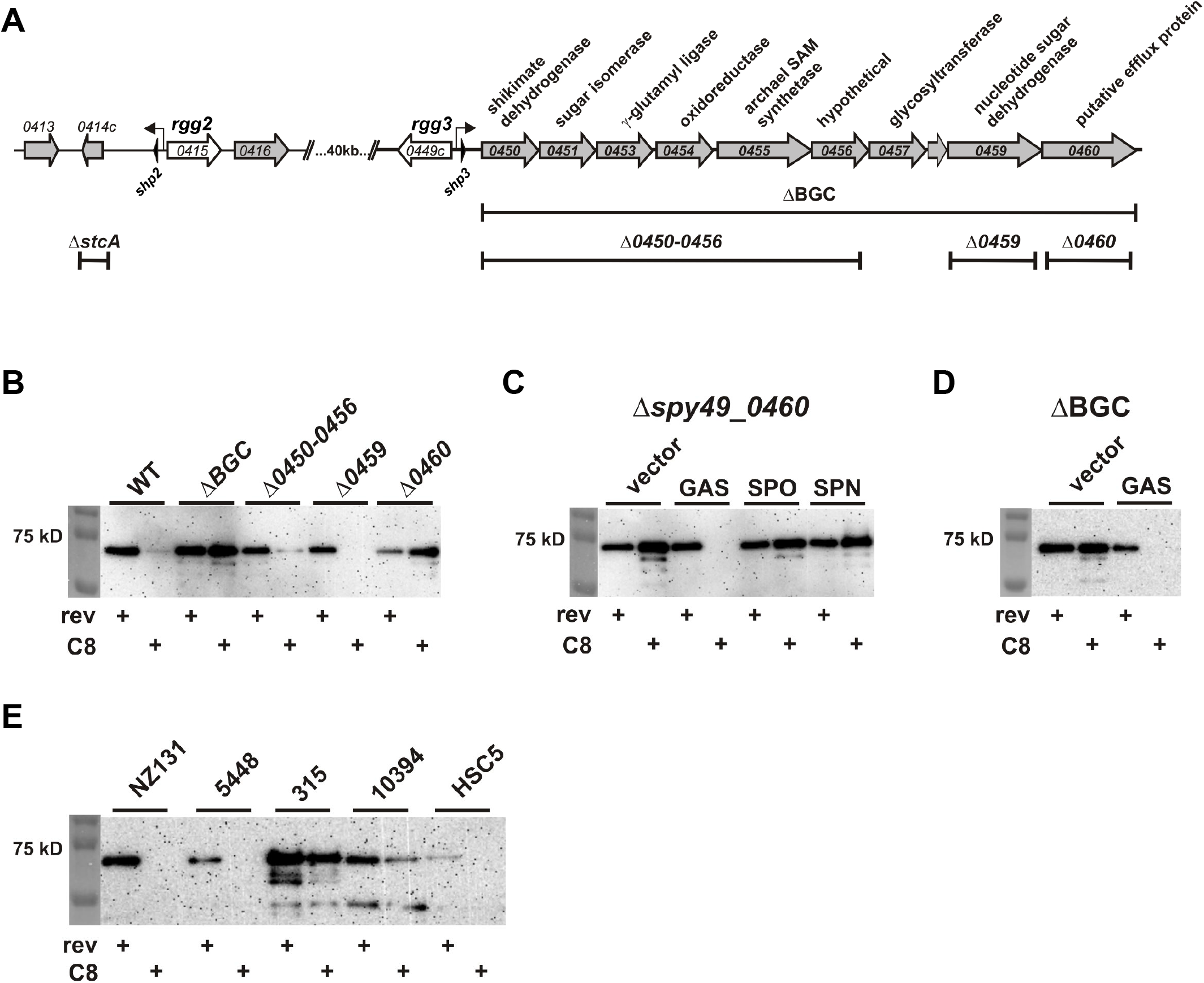
Reduction of streptolysin O in response to QS induction is dependent on a major facilitator superfamily protein, Spy49_0460. Supernatants from late log phase cultures treated with 100 nM reverse (rev) or C8 peptide were collected and western blotting with anti-SLO antisera was performed. **A**. Genetic loci regulated by Rgg2/3 QS. **B**. SLO accumulates in supernatants from mutants that lack *spy49_0460*. **C**. Complementation of Δ*spy49_0460* with MFS homologues from GAS, *S. porcinus* (SPO), or *S. pneumoniae* (SPN) expressed from the P_*shp3*_ promoter. **D**. Complementation of ΔBGC with GAS MFS protein *spy49_0460*. **E**. SLO in supernatants collected from additional WT GAS strains after induction of quorum sensing with C8 peptide: NZ131 (M49), 5448 (M1), MGAS315 (M3), MGAS10394 (M6), HSC5 (M14).

Rgg/SHP pairs are widely conserved in *Streptococcus* species, with some genomes containing multiple paralogs. A search for Rgg3 and SHP3 specifically, identified homologues in all *Streptococcus porcinus* genomes sequenced to date and in approximately one-third of *Streptococcus pneumoniae* genomes ^13^. In *S. porcinus, rgg3*/*shp3* lie next to a BGC containing homologues with 77-90% similarity to the proteins in the GAS BGC (Table S1); interestingly, *spy49_0458* in GAS appears to be a C-terminal remnant of what is annotated as an adenylation domain-containing nonribosomal peptide synthetase in *S. porcinus*. All but one of the *S. porcinus* genomes contain a homologue of Spy49_0460 and encode additional ABC transporter genes downstream of this gene. In the *S. pneumoniae* strains that contain *rgg3* and *shp3*, these genes are located next to one of four different BGCs – two encode predicted lantibiotic synthesis genes and do not have a Spy49_0460 homologue, and two contain biosynthetic genes unrelated to those of the GAS BGC but include a Spy49_0460 homologue. One of these latter pneumococcal Rgg3-SHP3 pairs has been shown to positively regulate surface polysaccharide expression, the overexpression of which decreases biofilm formation and in vivo fitness ^13^. The *S. porcinus* str. Jelinkova 176 and *S. pneumoniae* R6 homologues of Spy49_0460 are 88 and 89% similar to the GAS protein, respectively, thus we wondered whether these transporters could complement the GAS *spy49_0460* mutant. GAS, *S. porcinus*, and *S. pneumoniae* MFS transporter genes were cloned downstream of a P_*shp3*_ promoter, integrated into the GAS Δ*spy49_0460* genome and tested for their ability to complement the deletion strain for regulation of SLO production. Complementation of the *spy49_0460* mutant with the native gene added back in single copy restored the QS-responsive reduction in SLO (Fig. 2C). However, despite the degree of similarity between the MFS transporters (and the BGC itself in the case of *S. porcinus*) neither of the other species’ proteins were able to complement the GAS *spy49_0460* mutant. Surprisingly, introduction of *spy49_0460* complemented the entire BGC deletion (Fig. 2D) suggesting that the product of the BGC does not play a role in altered SLO production. Correspondingly, although *spy49_0460* appears at the distal end of the BGC transcript, this gene is not required for suppression of macrophage immune responses, the other phenotype linked with these genes ^6^. Thus, the MFS transporter appears to have a function independent of that of the rest of the BGC. Finally, we tested whether QS induction led to less SLO in other wild-type GAS serotypes. While basal levels of the toxin varied between strains, there was a trend for decreased SLO in all cells treated with SHP3-C8 (Fig. 2E).

### Deletion of *spy49_0460* leads to transcriptional changes for surface-associated virulence factors

To investigate the mechanism underlying the decrease in secreted SLO protein, we used luciferase (*lux*) reporter assays to monitor gene expression in response to the C8 pheromone. Our goals were to monitor changes in reporter activity in response to QS-induction (+C8) for each genotype and to compare changes in reporter activity in response to C8 peptide in the presence and absence of the MFS transporter. Streptolysin O is encoded in an operon downstream of the nicotinamide adenine dinucleotide (NAD)-glycohydrolase (*nga*) toxin and its immunity factor, *ifs*. The genomic sequence upstream of *nga* was cloned in front of bacterial luciferase genes, and the resulting reporter construct was introduced into wild-type and Δ*spy49_0460* strains. In wild-type bacteria, P_*nga*_-*lux* reporter activity was ∼6-fold lower at late log phase after treatment with 100 nM C8 peptide relative to the reverse peptide control (Fig. 3A). In contrast, treatment of Δ*spy49_0460* with C8 did not decrease P_*nga*_-*lux* activity to the same extent, and C8- and rev-treated cells exhibited similar reporter activity by late log phase. Luciferase reporter phenotypes of Δ*spy49_0460* could be complemented by the addition of *spy49_0460* on a plasmid (Fig. S2). These data are consistent with the results observed in the anti-SLO western blot (Fig. 2B) and suggest that transcriptional changes are responsible for the decrease in SLO protein and hemolytic activity (Fig. 1A) in QS-ON cells.

**Figure 3.**
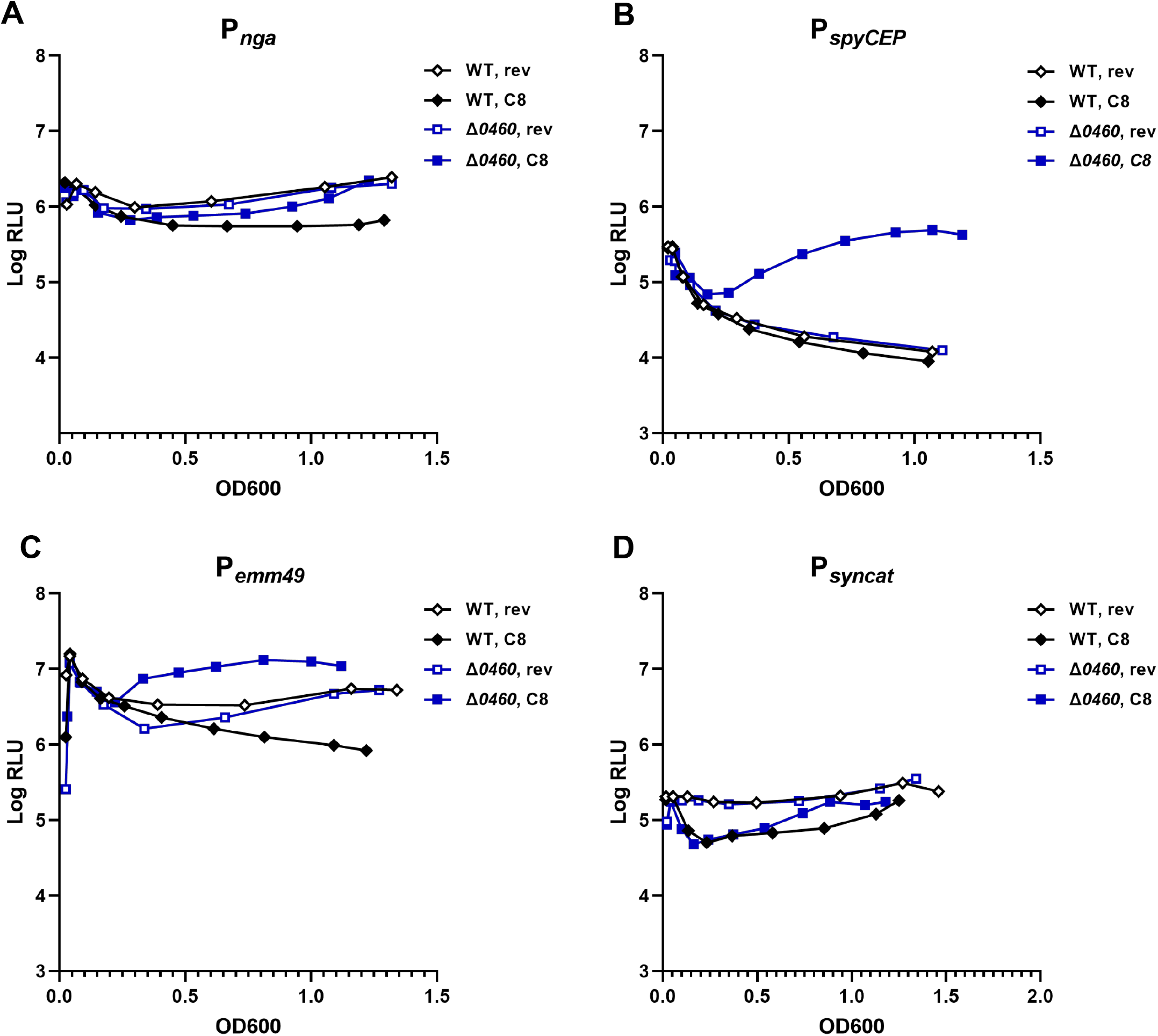
Transcriptional changes of several target genes in response to QS induction. WT and Δ*spy49_0460* strains containing luciferase reporters were grown in the presence of 100 nM reverse (rev) or C8 peptide and growth and light production were monitored. **A**. P_*nga*_ (streptolysin O). **B**. P_*spyCEP*_. **C**. P_*emm49*_ (M protein). **D**. P_*syncat*_ (constitutive). Data shown are representative of experiments performed at least three times.

We next examined the effect of QS induction on two additional secreted virulence factors, SpyCEP and M protein. SpyCEP (PrtS/ScpC) is a cell envelope serine protease that can cleave host IL-8 and interfere with recruitment of host neutrophils critical for controlling infection ^14^. M protein is a highly variable surface antigen mediating resistance to phagocytosis, adherence to host cells, and in some cases, invasion into host cells ^15, 16^. The genomic sequence upstream of each of these genes was cloned in front of luciferase genes, and the resulting reporter constructs were introduced into wild-type and Δ*spy49_0460* strains. The wild-type P_*spyCEP*_-*lux* reporter strain exhibited little difference in luciferase activity in response to C8 versus reverse peptide, and reporter activity in uninduced Δ*spy49_0460* was similar (Fig. 3B). Surprisingly, addition of C8 to Δ *spy49_0460* P_*spyCEP*_-*lux* led to a >40-fold increase in reporter activity relative to C8-treated wild-type and uninduced Δ*spy49_0460*. For P_*emm49*_-*lux*, addition of C8 peptide to wild-type led to a ∼6-fold decrease in reporter activity, similar to P_*nga*_-*lux* (Fig. 3C). In contrast, C8 treatment of Δ*spy49_0460* led to increased reporter activity relative to reverse peptide-treated cells, and ∼20-fold higher P_*emm49*_-*lux* activity relative to QS-ON wild-type. Finally, we measured the effect of QS induction on a constitutively-expressed synthetic promoter, P_*syncat*_. Addition of C8 peptide led to a modest (∼4-fold) decrease in reporter activity for both strains relative to the reverse peptide controls, but the deletion of *spy49_0460* had a limited effect on reporter activity overall (Fig. 3D).

To assess whether levels of SpyCEP and M protein correlated with our transcriptional reporter assays, late log culture supernatants were harvested and analyzed by western blot. As with SLO, induction of QS by the addition of C8 led to decreased levels of both SpyCEP and M protein produced by wild-type cells (Fig. 4). There was a greater decrease of SpyCEP (Fig. 4A) than we anticipated based on the luciferase reporter data (Fig. 3A), and we cannot rule out the role of proteolysis in protein turnover at this time. In contrast, SpyCEP and M protein levels in Δ*0460* culture supernatants were reflective of the increases we observed using the transcriptional reporter strains, with increased protein in C8-treated samples. We note that NZ131 (serotype M49) is phylogenetically classified as Clade X, with M proteins of this clade reported to bind the Fc portion of immunoglobulins ^17^. While we are unable to determine if the bands detected by the antisera used in Fig. 4B correlate to Fab- versus Fc-mediated binding, the loss of the bands in the Δ*emm49* mutant suggest that there is specificity for the M49 protein. From these data, we conclude that Rgg-SHP QS induction leads to both promoter-dependent and Spy49_0460-dependent changes in transcription that are largely reflected in the amount of protein present in the culture supernatant.

**Figure 4.**
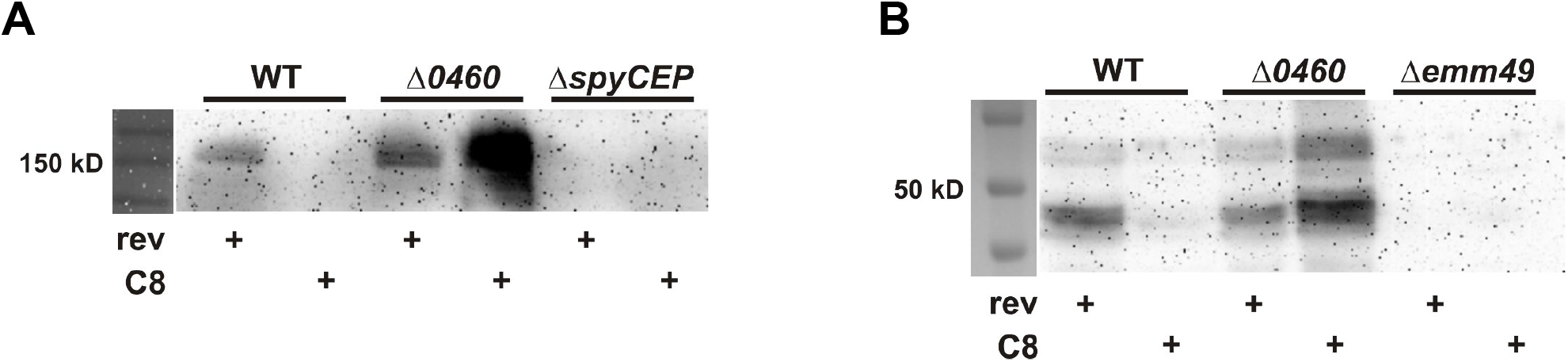
SpyCEP and M49 protein are reduced in a Spy49_0460-dependent manner in response to QS induction. Late log phase culture supernatants were analyzed by western blot using **A**. Anti-SpyCEP or **B**. Anti-M antisera.

### Spy49_0460 mediates QS-dependent sensitivity to aminoglycosides

In previous studies, we observed a SHP-dependent increase in cells’ sensitivity to aminoglycoside antibiotics kanamycin and streptomycin ^3^. Measuring the growth of rev- or C8-treated wild-type, *rgg2* (QS-OFF), *rgg3* (QS-ON) and *spy49_0460* mutants in increasing concentrations of kanamycin demonstrated that the MFS protein is responsible for this phenotype; deletion of the transporter resulted in the loss of C8-dependent drug sensitivity similar to the *rgg2* mutant (Fig. 5A). As with SLO production, this phenotype could be complemented with the GAS but not *S. porcinus* or *S. pneumoniae* alleles of the MFS transporter (Fig. 5B). We also observed that the expression of *spy49_0460* in the BCG mutant was sufficient to confer increased sensitivity to the drug, suggesting the rest of the operon does not play a role in sensitivity to the antibiotic (Fig. 5C). Finally, as a complementary experiment, we expressed GAS, *S. porcinus*, or *S. pneumoniae* MFS transporter genes in *S. porcinus* and observed that only the GAS *spy49_0460* led to increased sensitivity to kanamycin (Fig. 5D). Taken together, these data suggest that the increased sensitivity of QS-ON cells to aminoglycosides is due to increased expression of the MFS transporter and that the GAS protein has a higher affinity for this substrate than the other species’ transporters despite ∼89% overall similarity at the amino acid level.

**Figure 5.**
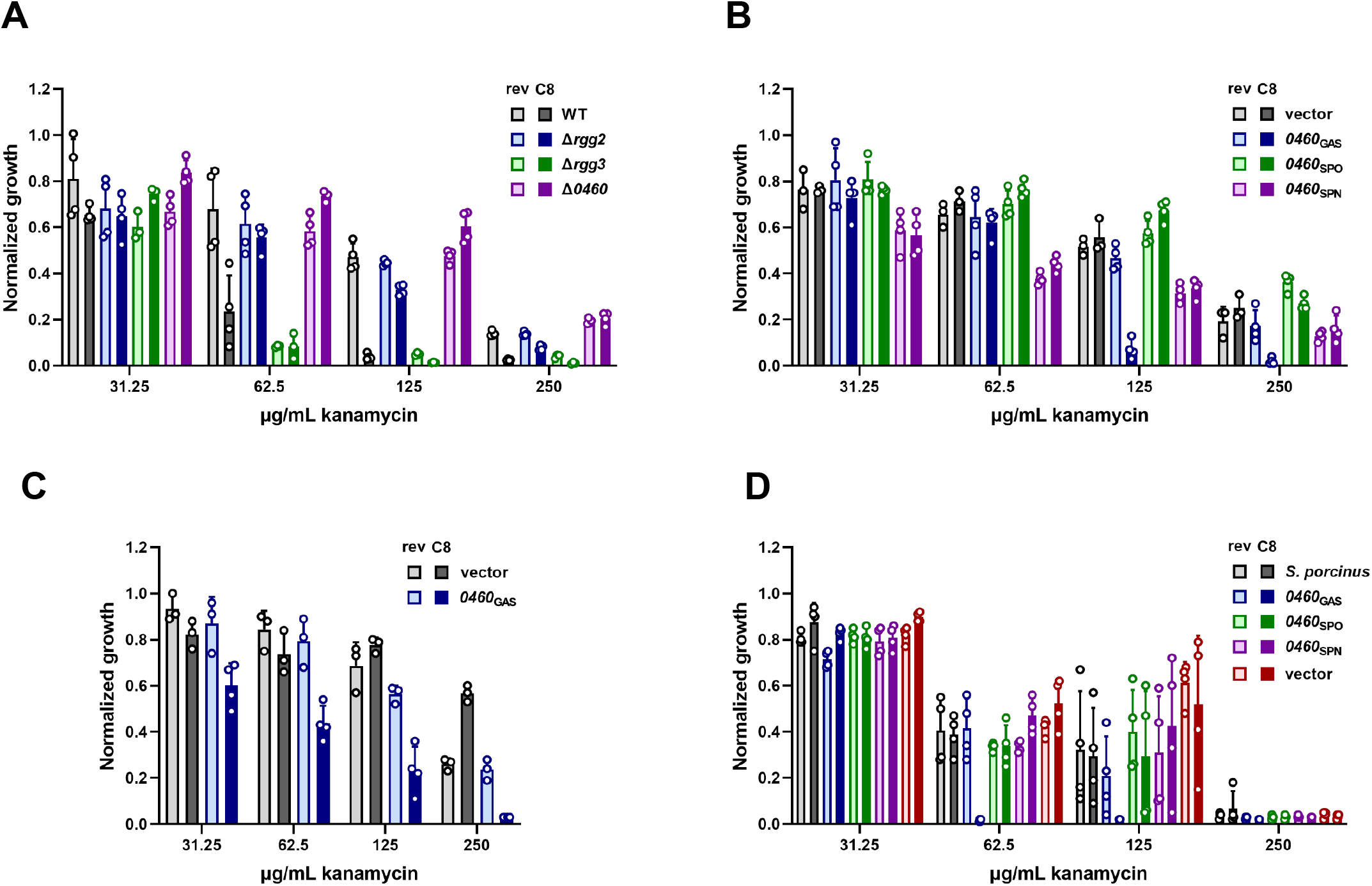
Expression of *spy49_0460* leads to increased aminoglycoside sensitivity. Kanamycin sensitivity was determined using a broth dilution assay in which mid log phase reverse (rev) or C8 peptide-treated bacteria were back diluted to a low OD600 in increasing concentrations of antibiotic. **A**. WT, *rgg2, rgg3* and *spy49_0460* mutants. **B**. Complementation of Δ*spy49_0460* with GAS, *S. porcinus* (SPO) or *S. pneumoniae* (SPN) MFS transporter genes expressed from a P_*shp3*_ promoter. **C**. Complementation of the ΔBGC mutant with P_*shp3*_*-spy49_0460*. **D**. Expression of GAS, *S. porcinus* (SPO) or *S. pneumoniae* (SPN) MFS transporter genes in *S. porcinus*. Data shown are mean and standard deviation from at least three biological replicates.

## Discussion

While we have made progress in understanding how streptococcal Rgg2/3 signaling is induced and works mechanistically, the full benefit this communication system confers to the bacteria and its relevance in infection remains unknown. Our current study identified a QS-regulated transporter, Spy49_0460, whose expression leads to decreased transcription of at least three secreted virulence proteins: the pore-forming cytolysin, streptolysin O; the chemokine-cleaving protease, SpyCEP; and the antiphagocytic surface antigen, M protein. In support of these observations, a recent proteomics study from our lab confirmed that streptolysin O and M49 are significantly less abundant in the secreted fraction of proteins from QS-induced cells ^18^. Spy49_0460 was previously identified in whole genome screens for genes important for virulence in a zebrafish model of necrotizing fasciitis ^19, 20^, survival in human blood ^21^ and pharyngitis in a nonhuman primate model ^9^, but the mechanisms underlying its importance in these models have not been determined.

Spy49_0460 is a member of the major facilitator superfamily of transporters, an extremely large and diverse contingent of proteins found in all kingdoms. These transporters use secondary active transport in which the transport of one substrate against its concentration gradient is coupled to the transport of another molecule or ion down its gradient. Substrates for these transporters include antibiotics, sugars, and metals among other small molecules, and it is not uncommon for transporters to have promiscuous transport capacity. While the use of this transport mechanism and the tendency of these proteins to have 12 transmembrane domains are hallmarks of this superfamily, primary amino acid sequence conservation is very low which makes it difficult to predict the precise functions of a given transporter. However, the role of MFS transporters from other bacterial pathogens in interactions with host cells has been documented ^22^.

Although annotated as a putative efflux permease or macrolide efflux pump, there is currently no evidence to suggest that expression of Spy49_0460 leads to increased antibiotic resistance. Surprisingly, deletion of *spy49_0460* increased resistance to the aminoglycoside kanamycin (Fig. 5). This correlates with a previous study where we observed that Rgg2/3 QS induction (which increases *spy49_0460* expression) also led to a 16-fold increase in sensitivity to streptomycin and a two-fold increase in sensitivity to both erythromycin and spectinomycin ^3^. A possible explanation for this observation, particularly with respect to kanamycin and streptomycin, is that aminoglycosides resemble the native cargo of this transporter (glycosidic, polycationic) and the promiscuity of the transporter allows increased drug influx when more copies are expressed. At least two fungal MFS transporters, Mdr1 and ScQdr2, have documented roles in both drug efflux and uptake ^23^, underscoring the challenge in assigning function to individual members of this large and diverse family.

A PHYRE search using the GAS amino acid sequence revealed structural matches to other MFS proteins including the human iron export protein ferroportin-1, the bacilysin/siderophore transporter BacE, and the lipoteichoic acid flippase LtaA ^24^. The former two hits are potentially interesting given that we previously observed that Rgg2/3 signaling is induced under conditions of iron limitation ^3^ and *spy49_0460* is located at the end of a BGC, and the latter is interesting due to the possibility that the BCG produces a factor that alters the GAS cell surface and its subsequent interactions with macrophages ^6^. However, we have not found a role for the BGC in iron uptake or a role for Spy49_0460 specifically in macrophage suppression to date.

At this time, we suggest the influence of Rgg2/3 on *slo, emm49* and *spyCEP* via Spy49_0460 is indirect. We showed previously that a 23-nt conserved Rgg binding sequence is present in only two locations in the GAS genome – directly upstream of *shp2*-*stcA* and *shp3*-*BGC*, the two loci directly regulated by the Rgg2 and Rgg3 transcription factors ^25^. An additional interesting observation was that the divergence in target gene transcription between wild-type and Δ*spy49_0460* strains did not become evident until ∼100 minutes after the addition of inducing pheromone (Fig. S3); this in contrast to the transcriptional response of P_*shp*_ promoters to C8 peptide which is detectable within minutes ^4^. This suggests that the downregulation of *slo, spyCEP*, and *emm49* transcription could be due to an accumulation of changes in cell physiology or stress that lead to the observed phenotype. In support of this idea, our recent proteomics study also detected nucleotide stress and a link to the stringent response in QS-ON cells ^18^. Alternatively, the Rgg2/3 regulon is induced within one hour of infection of the murine nasopharynx ^10^, though little is known about its expression later in infection. It is tempting to speculate that while SLO, SpyCEP and M protein play a critical role in the early events of infection, the bacteria might benefit from the ability to fine tune these genes’ expression via Rgg2/3 QS and Spy49_0460 as the infection progresses or when different conditions are encountered within the host.

## Methods

### Bacterial strains and media

GAS and *S. porcinus* were routinely cultured on Todd-Hewitt (TH; BD) + 2% yeast extract (Y; VWR) broth or agar. A chemically-defined medium (CDM; ^4, 26^) + 1% glucose was used for all assays. When appropriate, antibiotics were included at the following concentrations: chloramphenicol, 3 μg/mL; erythromycin, 1 μg/mL; kanamycin, 100 μg/mL, spectinomycin, 150 μg/mL. *E. coli* were routinely cultured on Luria broth or agar with antibiotics used at the following concentrations: chloramphenicol, 10 μg/mL; erythromycin, 500 μg/mL; kanamycin, 150 μg/mL; spectinomycin, 150 μg/mL.

### Construction of plasmids and mutant strains

All strains, plasmids and primers are described in Tables S2 and S3. Unless otherwise indicated, DNA from wild-type M49 GAS strain NZ131 was used as the template for PCR reactions. Deletions in the NZ131 parent strain were achieved by the introduction of deletion plasmids by electroporation followed a two-step, temperature-dependent selection process as described previously ^4, 27^. To construct the Δ*sagA* mutant, genomic regions flanking and including the start and stop codons of *sagA* were amplified and fused by PCR using primers JC71-74 and ligated into pFED760 to make pJC148. To construct the Δ*emm49* mutant, fragments encompassing the upstream flanking region, *aphA3* kanamycin resistance gene (pOsKar template) and downstream flanking region were amplified by PCR using primers RW137-RW142 and cloned into pFED760 using the NEBuilder HiFi Assembly method (New England Biolabs) to make pRVW47. To construct the complementation plasmids, MFS transporter homologue genes from NZ131 (*spy49_0460*; primers JC590/591), *S. pneumoniae* CP2000 (*spr0971*; primers JC603/604) or *S. porcinus* str. Jelinkova (*strpo_0488*; primers JC612/613) were amplified from the indicated template and cloned downstream of a P_*shp3*_ promoter fragment (primers JC131/JC592) in pLZ12-Sp (pJC417) or p7INT (pJC420, pJC430, pJC434). To construct reporter plasmids, promoter regions for *emm49* (primers JC94/JC108; pJC167), the synthetic *cat* cassette (pEVP3 template; primers JC208/JC412; pJC318), or *nga* (primers RW232/RW233; pRVW71) were amplified and cloned in front of bacterial luciferase genes.

### Starter culture preparation

Starter cultures were prepared by culturing single colonies of GAS in THY to an OD600 of ∼0.7. Glycerol was added to a final concentration of 20%, then individual aliquots were frozen and stored at −80° C. For all experiments, aliquots were thawed, and the bacteria were pelleted and resuspended in CDM before dilution into fresh CDM.

### Synthetic peptides

Previous studies identified the C-terminal eight amino acids of SHP2 and SHP3 as the minimum peptide length with activity ^28^. SHP2-C8 and SHP3-C8 initiate transcription at both the *shp2* and *shp3* promoters and thus are functionally equivalent. Crude preparations of synthetic SHP3-C8 (C8, DIIIIVGG) or SHP3-C8 reverse (rev, GGVIIIID) were obtained from Genscript or Neo Peptide, respectively, reconstituted to 1 mM in DMSO and stored at −80°C. Working stocks at 100 μM were prepared in DMSO and stored at −20° C.

### Red blood cell hemolysis assay

Supernatants were harvested from cultures grown to late-log phase (OD600 = 1.0) in CDM, and dithiothreitol (DTT) was added to a final concentration of 4 mM; where indicated, water-soluble cholesterol (Sigma) was added to a final concentration of 2 mg/mL. The supernatants were incubated at room temperature for 10 min then serially diluted in phosphate-buffered saline (PBS) + 4 mM DTT in a microtiter plate. Sheep red blood cells (RBC; Cocalico Biologicals) were washed 2-3 times with cold PBS and diluted to a final concentration of 10% in PBS. 25 μL of washed RBC were added to 100 μL of diluted supernatant followed by incubation at 37° C for 30 min. Sterile water, PBS, and CDM were included as the positive, negative and medium controls for hemolysis, respectively. Following incubation, the plates were spun at ∼800 x g for 5 min, then 75 μL of the supernatant was transferred to a fresh plate and the OD450 nm was recorded. Absorbance values were normalized to the 100% lysis control.

### Western blotting

For SLO, supernatants from cultures grown in CDM to late-log phase (OD600 = 1.0 for NZ131 and isogenic mutants; OD600∼0.5-0.7 for other wild-type strains) were concentrated by precipitation with 10% trichloroacetic acid (Sigma). For SpyCEP and M protein, cultures were grown to OD600 = 0.8, then cells were pelleted and lysed with PlyC ^29^. After separation by SDS-PAGE, proteins were transferred to a PVDF membrane (Biorad) and blocked with 4% bovine serum albumin + 1% nonfat milk in Tris-buffered saline + 0.05% Tween 20. Incubation with primary antibodies or antisera was performed overnight at 4° C with dilutions as follows: anti-SLO (Abcam), 1:6,000; anti-SpyCEP (gift from S. Sriskandan), 1:1,000; anti-M22 (gift from G. Lindahl) ^30^, pre-adsorbed against heat-killed Δ*emm49* then diluted to 1:1,000. Note: the anti-M22 antisera was predicted to recognize M49 on the basis of similar C repeats (G. Lindahl, personal communication). A horseradish peroxidase-conjugated anti-rabbit secondary antibody (Biorad) was used at a dilution of 1:20,000, and chemiluminescence was detected using Supersignal West Femto substrate (Pierce) and a FluorChem E imager (ProteinSimple). To confirm equivalent amounts of sample were loaded on each gel, the PVDF membranes were subsequently stained with India ink (Remel; ^31^) and photographed (Fig. S4).

### Luciferase assays

Starter cultures of strains containing the luciferase reporters for the indicated genes were thawed, pelleted, then resuspended and diluted into fresh CDM. At t=0 and subsequent time points, the OD600 was recorded, then 50 μL of culture was transferred to an opaque white plate, exposed to decyl aldehyde (Sigma) fumes, and light production was recorded. Counts per second were normalized to OD600 to obtain relative light units (RLU). Synthetic SHP3-C8 or reverse peptide was added to a final concentration of 100 nM after 60 min of growth. Although P_*nga*_ and P_*spyCEP*_ reporter plasmids contain *luxCDE* genes capable of producing endogenous substrate, we found that the amount produced was limiting, thus exogenous decyl aldehyde was also added in assays using these reporters.

### Aminoglycoside MICs

Assays were performed similarly to those described previously ^3^. Briefly, starter cultures were inoculated into CDM and pre-treated with C8 or reverse peptide for 1 hour before back dilution into microtiter plates containing 2-fold serial dilutions of kanamycin and 100 nM of peptide in a final volume of 150 μL. Plates were parafilmed and incubated overnight at 37° C in 5% CO_2_. After 24 h, cells were gently resuspended, and 75 μL from each well was transferred to a fresh plate. The absorbance at 600 nm was recorded, and growth at each concentration of antibiotic was normalized to the no drug control.

## Acknowledgements

We thank Drs. Shiranee Sriskandan and Gunnar Lindahl for their kind gifts of anti-SpyCEP and anti-M22 antisera, respectively; Dr. Bobbi Xayarath for technical recommendations for western blotting; and Dr. Artemis Gogos for technical help.

